# Efficiently Quantifying DNA Methylation for Bulk- and Single-cell Bisulfite Data

**DOI:** 10.1101/2023.01.27.525734

**Authors:** Jonas Fischer, Marcel H. Schulz

## Abstract

**Motivation:** DNA CpG methylation (CpGm) has proven to be a crucial epigenetic factor in the gene regulatory system. Assessment of DNA CpG methylation values via whole-genome bisulfite sequencing (WGBS) is, however, computationally extremely demanding.

**Results:** We present FAst MEthylation calling (FAME), the first approach to quantify CpGm values directly from bulk or single-cell WGBS reads without intermediate output files. FAME is very fast but as accurate as standard methods, which first produce BS alignment files before computing CpGm values. We present experiments on bulk and single-cell bisulfite datasets in which we show that data analysis can be significantly sped-up and help addressing the current WGBS analysis bottleneck for large-scale datasets without compromising accuracy.

**Availability:** An implementation of FAME is open source and licensed under GPL-3.0 at https://github.com/FischerJo/FAME.

## Introduction

DNA methylation has been shown to be an essential building block of the gene regulatory system (Mattei, Bailly, and Meissner 2022), being often referenced as the “fifth base”. As an epigenetic mark, it has been shown to be crucial for dynamic gene regulation throughout development and aging (Smith and Meissner 2013; Bashkeel et al. 2019; Greenberg and Bourc’his 2019; Field et al. 2018) and aberrant methylation has been reported in different diseases, especially having a broad impact on cancer (Lister and Ecker 2009; Kulis and Esteller 2010; Papanicolau-Sengos and Aldape 2022; Huang et al. 2014; Wang et al. 2022). DNA methylation can be measured large scale using methylation arrays (Pidsley et al. 2016) or using sequencing (Meissner et al. 2005; Krueger et al. 2012). Assessment of genome-wide DNA methylation status via bisulfite sequencing (BS) is considered as the gold standard for methylation profiling. The current developments in sequencing technology, however, led to a drastic increase in the amount of data produced, on the one hand due to higher throughput and on the other hand due to the development of single-cell BS approaches in which hundreds of individual cells are analyzed (Schwartzman and Tanay 2015; Smallwood et al. 2014; Farlik et al. 2015; Liu et al. 2021).

While there have been many attempts to speed-up the computation of DNA CpG methylation (CpGm) values, all previous approaches have relied on a two-step procedure. Bisulfite reads are first aligned to the genome and then CpGm values are computed from read alignment files.

This separation into two steps creates an unnecessary overhead if the primary interest lies in the CpGm values, because bisulfite alignment files consume large amounts of disk space, imposing extra I/O time and disk resources for the computations. Furthermore, most methods are built around existing read alignment software that was originally developed for traditional DNA read alignment. However, WGBS reads are more difficult to align as read T nucleotides may map to genomic C nucleotides. This not only requires more specialized index structures compared to traditional DNA alignment (Supp. Fig. 1), but also introduces mapping ambiguity for read T nucleotides, referred to as the asymmetric mapping problem (Lister and Ecker 2009).

Based on how the asymmetric mapping problem is handled, available methods for WGBS read alignment can be classified into reduced alphabet (RA) and bisulfite-aware (BA) mapping approaches. RA mappers exchange C residues to T in all sequences (reads and reference), obtaining a reduced three letter alphabet, such that bisulfite converted cytosines do not count as sequencing errors during mapping. This reduction leads to false positive mappings in the genome (Fig. 1a) and, hence, less sensitive methylation calls. Popular RA mappers are Bismark (Krueger and Andrews 2011), BSSeeker (Chen, Cokus, and Pellegrini 2010; Guo et al. 2013), BratNova (Harris, Ounit, and Lonardi 2016), as well as gemBS (Merkel et al. 2019), and bwameth (Pedersen et al. 2014), which use efficient DNA aligners such as Bowtie (Langmead and Salzberg 2012) or bwa (Li and Durbin 2009) to build an index of CtoT and GtoA converted genome sequences.

**Figure 1:**
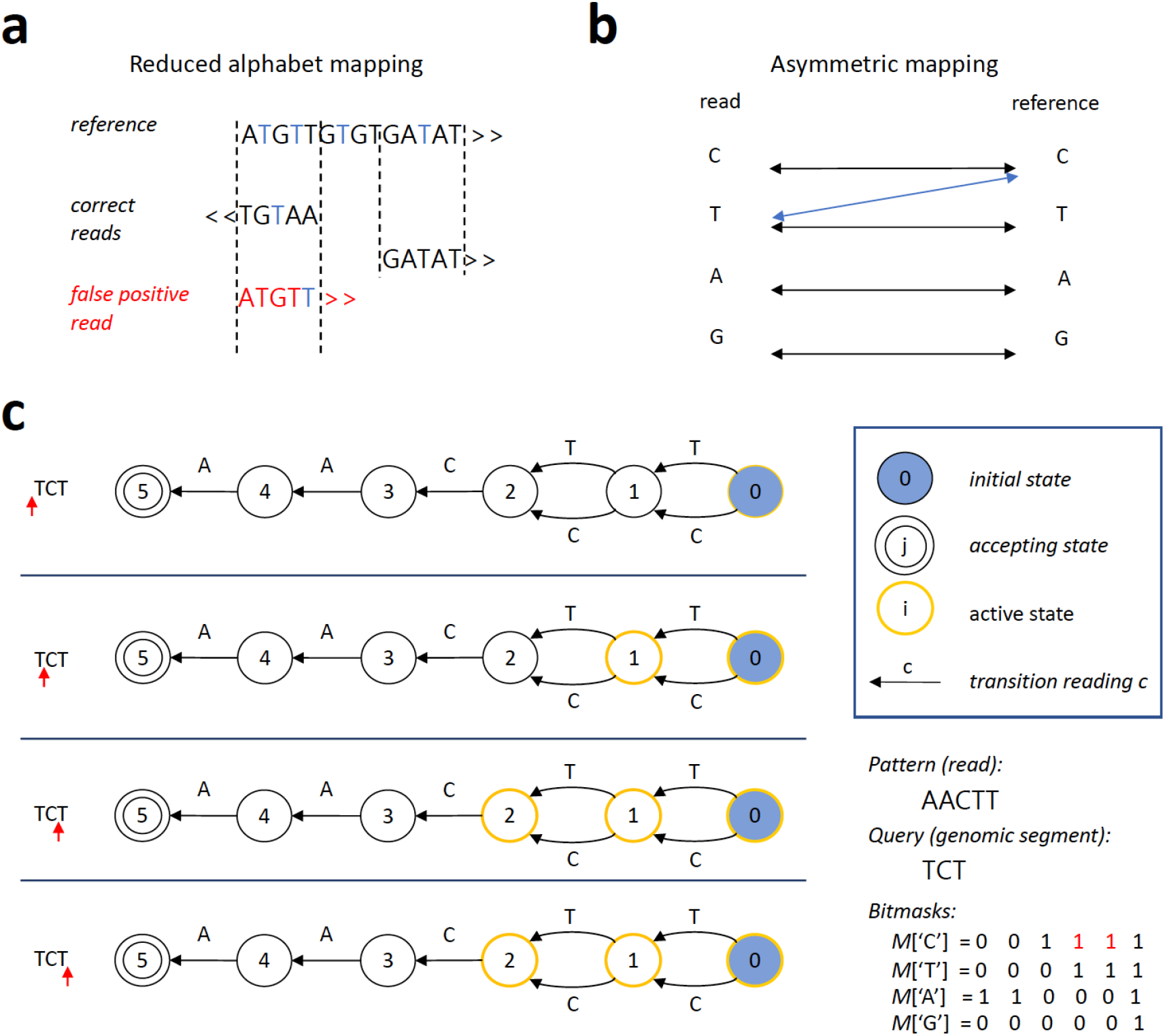
Alignment aware mapping. **a** Errors made through traditional reduced alphabet mapping. Former C reduced to T are given in blue, strand origin indicated by arrows. **b** Principle of asymmetric aware mapping, T can be mapped unidirectionally to C. **c** Example of asymmetric (bisulfite-aware) Shift-And automaton. Visualization of the query of the letter sequence (i.e., genomic segment “TCT” to the for the pattern “AACTT” (i.e., read). The current position in the segment (left side) is indicated by a red arrow.

While widely deployed, RA mappers do not correctly address the asymmetric mapping problem and have been shown to have reduced accuracy (Otto, Stadler, and Hoffmann 2012; Frith, Mori, and Asai 2012). This is because they do not properly account for wrong mapping positions due to the reduced alphabet. In contrast, BA methods use different approaches to resolve the mapping ambiguity: (i) realign potential reads at positions obtained with the reduced alphabet using a BA alignment method (Huang, Huang, and Chen 2018), (ii) use a *k*-mer based index with bitmasks to address converted cytosines (Xi and Li 2009), or (iii) directly solve the asymmetric mapping problem with a modified alignment algorithm (Otto, Stadler, and Hoffmann 2012; Frith, Mori, and Asai 2012). BA mappers such as Segemehl (Otto, Stadler, and Hoffmann 2012) are among the most accurate methods, but are very slow and, hence, hardly scale to the massive amounts of data currently being produced. While there have been many attempts to speed-up the bisulfite alignment process, the extended throughput in current single cell datasets is making analysis even more challenging and customized single cell analysis methods are needed (Wu et al. 2019).

Here we present FAME (Fast and Accurate MEthylation calling), the first BA mapping method with an index that is tailored for the alignment of BS reads with direct computation of CpGm values. Our method is built on a novel data structure that exploits gapped *k*-mer counting within short segments of the genome to quickly reduce the genomic search space. FAME is optimized to handle large single cell datasets. With the resulting lightweight index structure, FAME enables ultra-fast and parallel querying of reads without I/O overhead. We evaluated the performance of FAME on both synthetic and real data in comparison to the state-of-the-art tools. All experiments showcase the unique ability of FAME to retrieve as accurate results as the best state-of-the-art methylation caller in just a fraction of time. With orders of magnitude faster processing time and no need to write intermediate files to disk, it overcomes the current bottleneck in BS-seq processing, easily scaling to large single cell processing tasks.

## Approach

To solve bisulfite-aware WGBS mapping efficiently, we propose **F**ast and **A**ccurate **ME**thylation calling (FAME). Our method consists of two main building blocks, an index structure specifically tailored to model CpGm methylations, and an efficient alignment method that implements asymmetric mapping of read Ts to reference Cs based on the index. We next give a more detailed overview of FAME and then discuss the index and alignment separately.

### FAME

FAME is a bisulfite aware mapper that is built on top of a custom indexing and alignment method, an overview of which is given in Fig. 2. It leverages a gapped *k*-mer based index structure (Fig. 2a), that enables ultra-fast retrieval of small genomic segments of large genomes, where each of these segments contains a potential match for the given read. This index structure is fixed for a given reference genome and thus can be queried in parallel for multiple reads. Furthermore, it contains counting structures for all CpGs that allow a direct update of methylation rates – the measure of interest – and thus avoids expensive hard disk writes of intermediate alignment files. Using this index, FAME aligns a read in three phases, (i) MCpG candidate retrieval, (ii) verification of candidate MCpGs, and (iii) methylation calling (Fig. 2b). In the following, we explain the details of the index and how to use it to align a read and call methylation rates.

**Figure 2:**
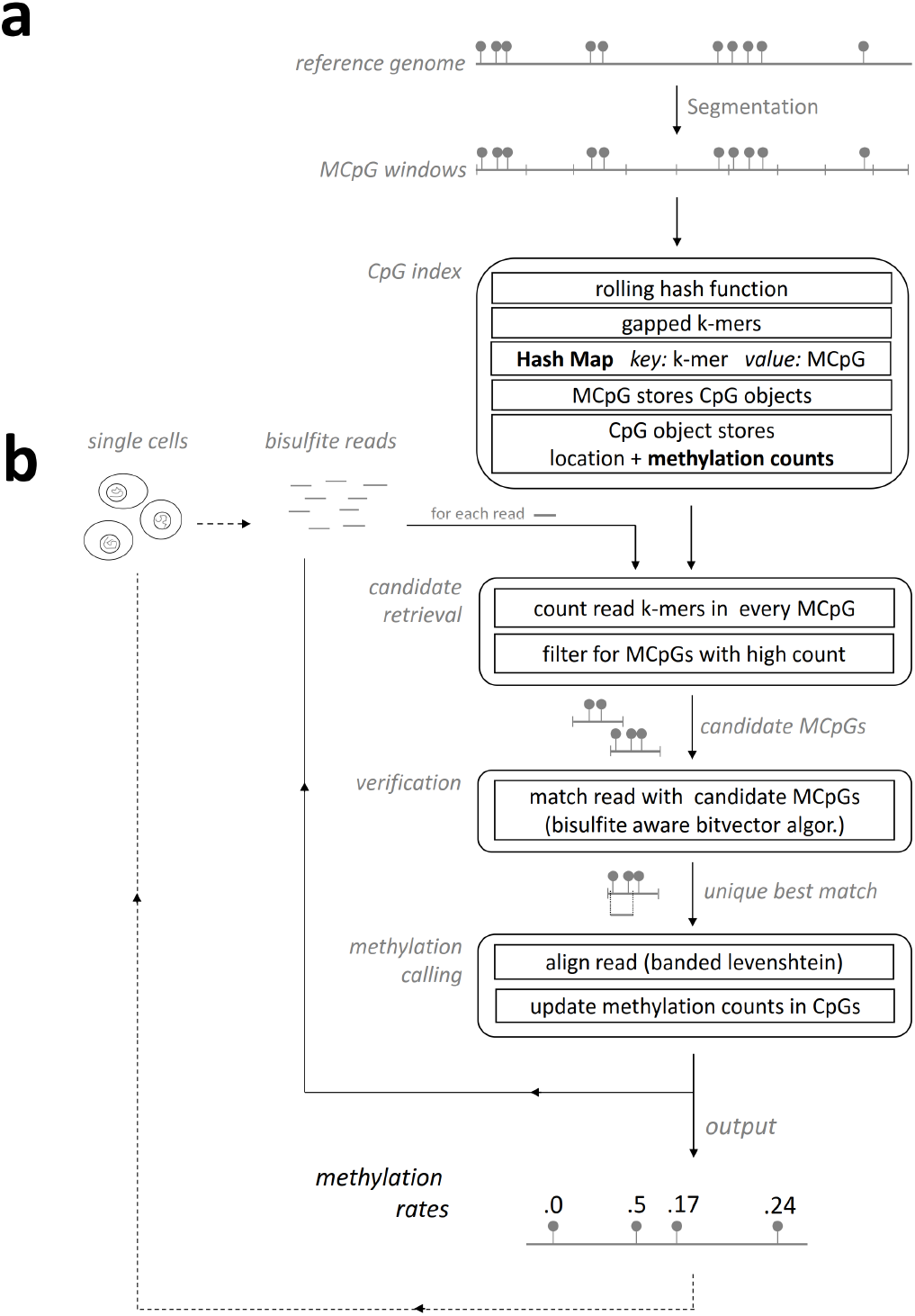
FAME workflow. General workflow of FAME for index construction (**a**) and read alignment (**b**). **a** For a given reference genome (top), a CpG index is constructed for all MCpG windows using a rolling hash function for gapped *k*-mers. **b** WGBS reads are searched with three phases: candidate retrieval, verification and methylation calling. Methylation rates are updated in a separat data structure, directly yielding methylation values without any realignment (bottom). FAME can process bulk or single cell datasets.

### Index

For index construction, a given reference genome is segmented into small windows of about 2kb length, called Meta CpGs (MCpGs). Each MCpG stores the set of CpGs it comprises and their respective genomic location. Every second *k*-mer within each MCpG is indexed using a fast rolling hash function from the library ntHash (Mohamadi et al. 2016), replacing all Cs by Ts in each spaced *k*-mer. All false positives introduced by this replacement are resolved in the alignment phase. Such spaced *k*-mers are substrings of length k that are masked with a spaced seed that consists of *k* positions that are either the original letter or a wildcard. A spaced *k*-mer thus matches a length *k* genomic sequence if for each position the letter in the *k*-mer and genomic sequence are equal, or the *k*-mer contains a wildcard. The hash values for spaced *k*-mers present in the reference genome, which are computed by ntHash, are used to populate a large hash table^1^, mapping spaced *k*-mers to the MetaCpGs they appear in.

To make the index more efficient, three optimizations are applied to the index. The first optimization, which is loss-less, removes all except one *k*-mer object belonging to the same MCpG within a hash table cell (i.e., *k*-mers that have the same hash value), as this information is redundant. The second optimization throws out a distinct *k*-mer that occurs more than a predefined threshold, as such *k*-mers are likely to belong to repetitive regions, which are problematic during alignment. The third optimization introduces a bitmask for all reference *k*-mers. This bitmask has a 1 for all positions with a T, 0 otherwise. For a given read *k*-mer, we can thus discard reference *k*-mers if the read *k*-mer has a C at any positions with a 1 in the bitmask. This allows to reduce the number of false positives by a quick bitmask check already in the candidate retrieval phase.

With the index structure at hand, we can now efficiently retrieve candidate regions for bisulfite-aware alignment, which is explained in the following section.

### Alignment

To align a read to the reference, FAME runs trough three phases (compare Fig. 2b). In the first phase, the *candidate retrieval*, each spaced *k*-mer of the read is looked up in the index using its hash value provided by ntHash. For each *k*-mer we, hence, get a collection of MCpGs where this *k*-mer might occur in. This way we obtain for each read an upper bound on how many of its *k*-mers are present in each MCpG. By setting a minimal number of k-mer occurrences *q* in a MCpG, most MCpGs can be excluded from further alignment steps.

In the second phase, similar to previous work (Otto, Stadler, and Hoffmann 2012), we employ a bisulfite-aware bitvector algorithm to solve the asymmetric mapping problem using edit distance and verify read matches to all candidate windows. In particular, FAME builds a Shift-And automaton for the read (Baeza-Yates and Gonnet 1992), but initializing the bitmask for Cs to all positions where a C **or a T** is present in the read. An example of a bisulfite aware Shift-And automaton is given in Fig. 1c, which can be easily generalized to an approximate Shift-And automaton (Supp. Fig. 2). All candidate MCpGs are queried to the automaton to find true matches for a given read.

For paired end reads the three steps work analogously, but are applied to both reads in parallel. An additional pruning step can be carried out after the first step by throwing out all MCpGs of one read that do not have a corresponding MCpG that lies within insert size distance away to the paired read.

In the third phase, the unique best hit – if one exists – is used to align the read to the original reference genome at this position and the CpGm counts are updated immediately. FAME applies a Dynamic Programming computation of the Levenshtein distance to find the best alignment between read and the matching subsequence of the reference leveraging banded alignment matrices, with the band width given by the number of errors in the unique best hit.

Storing all data in memory allows highly parallel implementation of all three steps and results in fast processing, while the memory consumption is similar to other methods, requiring less memory than Bismark or Segemehl (see Supp. Tab. 1). Further, there is no need for large disk storage capacities for intermediate files, a bottleneck imposed by other methods, which generate up to terabytes of data for a single single-cell dataset.

All alignment phases can also naturally be extended to support single cell experiments (compare Fig. 2). For that, reads are processed as batches, where each batch represents the read yield of a single cell. Each of these batches runs through the three alignment phases of FAME and methylation rates are estimated per batch and streamed into a file. This results in a matrix containing all single cell experiments as rows, avoiding any overhead caused by repeated loading of the index for each cell, or excessive I/O caused by writing out methylation calls for all CpGs for each individual cell, which usually have sparse coverage.

## Methods

In the following, we discuss data generation and preprocessing along with performance metrics used for our evaluation and comparison.

### Synthetic data preparation

To carry out an evaluation on data with known ground truth, we generate synthetic data resembling high quality WGBS reads. Each synthetic data set consists of 25 million 100bp reads sampled uniformly from chromosome 22 of human reference genome hg19. For each CpG in the genome we flip a coin and draw a methylation rate *p* either from *N* (0.2, 0.08) or *N* (0.8, 0.08), reflecting hypo- or hyper-methylated CpGs by Gaussians with low or high mean, respectively, as usually observed in bulk sequencing data. For each generated read, we methylate each CpG in the read with probability *p* given by the reference CpG. Conversion success rate is set to 99%, which is commonly observed in real WGBS experiments. Cytosines in non-CpG context are methylated with probability 0.01 reflecting real mammalian genomes. Finally, we introduced *l* errors, where *l* was drawn from a Poisson with *λ* = 0.5 to simulate base calling errors. For paired end reads we generated 25 million pairs of two reads generated as explained above, with the restriction that reads should only be between 100bp and 400bp apart from each other, drawn uniformly at random, reflecting the allowed insert size of paired end protocols. As we know the ground truth mapping positions of each read and the methylation counts for each CpG, we can compute the Root Mean Squared Error (RMSE) and Spearman correlation coefficient using all CpGm values. We considered all CpGs, where at least one tool mapped more than five reads. We measured the runtime as the sum of time by alignment and by methylation calling.

### WGBS and EPIC bead data processing

In preparation for the comparison on real data, we downloaded the 5 WGBS paired-read datasets of LNCaP cell by (Pidsley et al. 2016), along with corresponding EPIC bead methylation calls. We then processed the EPIC bead data using RnBeads (Assenov et al. 2014; Müller et al. 2019) to obtain methylation rates for all CpG positions on the EPIC bead array. For the two replicates, we averaged the methylation rates per CpG. The 5 WGBS read sets were pooled, leading to about 437 million reads. Adapters were trimmed using trim galore^2^, following the original approach by (Pidsley et al. 2016):

~~~
1 trim_galore --length 85 --max_n 1
     --paired --gzip
2 LNCaP_pooled_1.fastq.gz
     LNCaP_pooled_2.fastq.gz -o
     trimmed/
~~~

For the evaluation, we considered the 841, 708 CpG positions out of 843, 385 CpGs present on the array, where at least one tool mapped more than five WGBS reads. We considered the EPIC derived CpGm values as a baseline for this comparison.

### Single-cell WGBS data processing

To test available methods on a recent single-cell dataset, we consider data of induced pluripotent stem cells generated in a study by (Linker et al. 2019). Analogue to the original study, we preprocessed the single cell data using trim-galore.

~~~
1 trim_galore --paired --clip_R1 6
     --clip_R2 6 --gzip --length 90
2 CELLID_1.fastq.gz CELLID_2.fastq.gz
~~~

### Performance metrics

In previous work, the number of aligned reads has often been considered a good proxy for mapper performance. However, this measure is misleading as reads aligned to the wrong position are counted positively towards performance. As an extreme example, consider a mapper that maps all reads to the first position of chromosome 1, resulting in 100% aligned reads, which is of course a wrong alignment. Thus a better approach is to report the number of correctly mapped reads, doing an extensive postprocessing of alignments to compare to known ground truth read positions on synthetic datasets. For real data, such a ground truth is impossible to obtain.

With the actual goal of quantifying DNA methylation, we can mitigate this issue by evaluating how close the predicted methylation rate is to the actual methylation rate. Hence, the performance is good if and only if the alignment is good (i.e., correct). We can leverage orthogonal experiments measuring CpG methylation on real data, such as EPIC arrays, to obtain alignment independent methylation rates against which we can evaluate all WGBS estimates. We measure predicted against actual methylation rate in terms of the Root Mean Squared Error (RMSE):

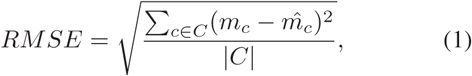

where *C* denotes the set of all CpGs with measurements and |*C*| denotes the size of this set. *m*_*c*_ denotes the ground truth (simulations or EPIC array) CpGm value of CpG *c* and 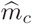 denotes the estimated value of one of the investigated tools based on WGBS data. Similarly, Spearman rank-based correlation was computed to measure the agreement between baseline CpGm values and estimates from the compared tools.

We consider all CpG sites where at least one tool mapped more than 5 reads (considering this CpG as mappable) and set the predicted methylation rate of tools that do not have any mapped reads at this CpG to 0 if the ground truth methylation rate is larger than 0.5, and to 1 if the ground truth methylation rate is smaller than 0.5. This introduces a small penalty if a tool is not able to cover a region during alignment, although it was possible.

## Results

We have developed FAME as a novel method for the quantification of DNA methylation rates at CpGs directly from bisulfite read data without producing intermediate alignment files (Fig. 2). We compare FAME against existing RA and BA mapping approaches in terms of runtime and quality of called methylation rates both on synthetic and real data. We note that there are a plethora of tools available and we have selected a competitive subset of those for our comparison, which are the most state-of-the-art algorithms that have full capabilities for alignment and methylation calling (more details, including why certain tools are excluded, in Supp. Tab. 2). The competing RA mappers were BratNova (Harris, Ounit, and Lonardi 2016), Bismark (Krueger and Andrews 2011), and gemBS (Merkel et al. 2019), the competing BA mappers were BSmap (Xi and Li 2009), and Segemehl (Otto, Stadler, and Hoffmann 2012) (method parameters in Supp. Sec. 5).

We measure the quality of called methylation rates in terms of Spearman correlation coefficient and the RMSE between predicted and baseline methylation rates. With ground truth methylation rates known for synthetic data, and methylation estimates available from orthogonal experiments for real data, we can thus evaluate each method’s ability to call methylation rates. Furthermore, methylation calls serve as a proxy for read alignment performance, as the methylation calls are good if and only if the alignments are good. Thus, in our evaluation we are not concerned with the issues of typical read alignment evaluations, where ground truth read positions are inherently hard to obtain for real data.

### Hyperparameter optimization

We first designed a simulation study to determine FAME’s default parameters on an independent synthetic dataset following the **in silico** data generation described in Section. The results for varying filter threshold *t* – the maximum times a *k*-mer is allowed to occur in the genome to be considered for the index – and the minimum number of reads *q* for a MCpG to be considered for alignment are given in Fig. 3. As expected, we can see that with larger *t* and smaller *q* the performance worsens but the runtime improves. These parameters provide a natural tradeoff between accuracy and runtime. We settle for default values *t* = 1500, *q* = 5, as they show a good balance between runtime and prediction accuracy and use them throughout the rest of the paper.

**Figure 3:**
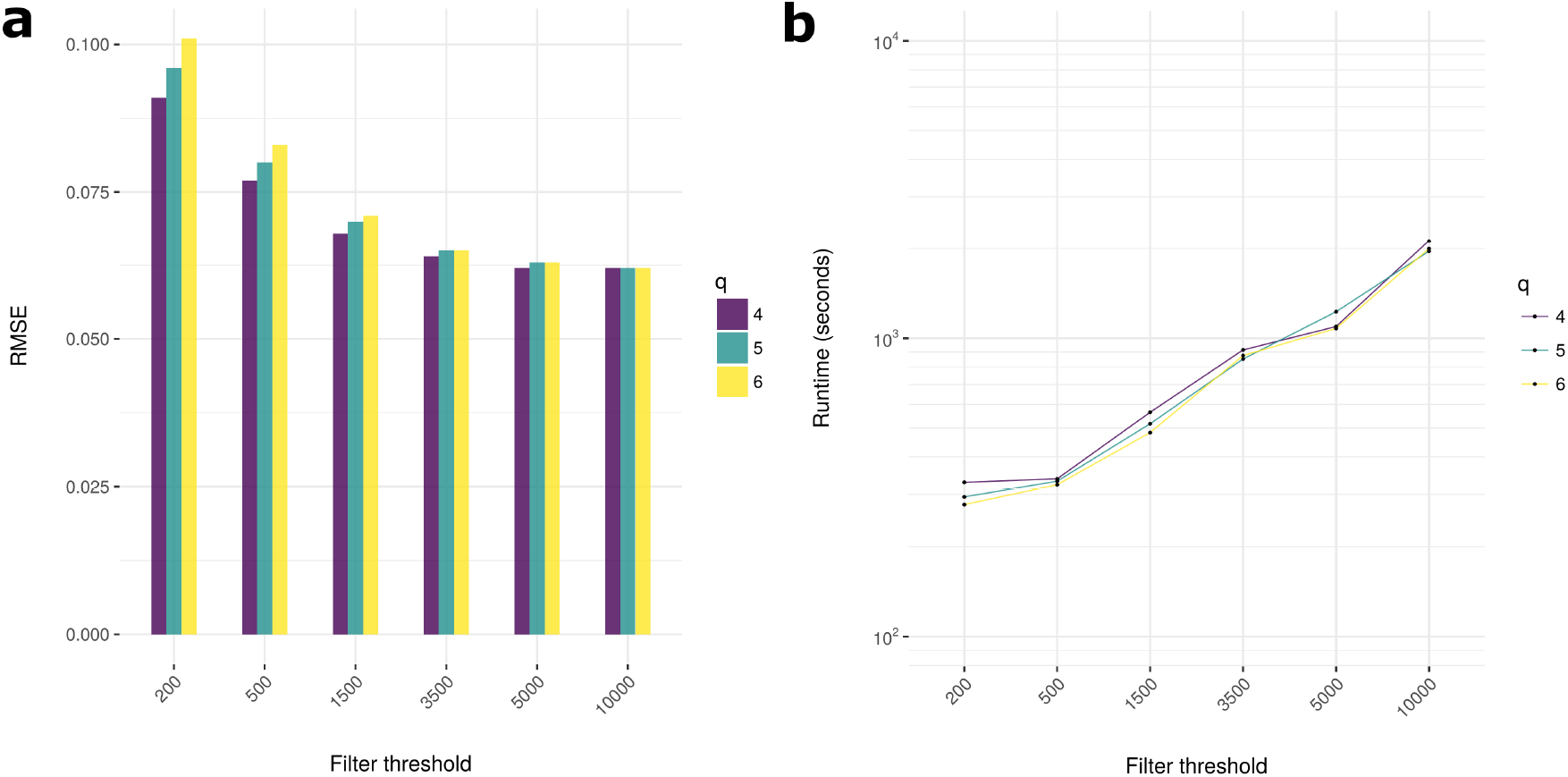
Results for different index parameters. Depicted are the results of a grid search for varying *q* (color), and filter threshold *t* (x-axis). (**a**) Root Mean Squared Error (RMSE, smaller is better) compared between actual and predicted methylation rates on the synthetic grid data set and (**b**) runtime for the same.

### DNA methylation calling on synthetic read set

Next, we generated synthetic data (25 million reads) sampled from human chromosome 22 resembling WGBS sequencing data (see Section). The performance of each tool was measured in terms of the combined runtime of read alignment and methylation calling on the whole genome, as well as quality of methylation calls. The results are given in Tab. 1 and Fig. 4a, a more detailed visualization of the called methylation rates can be found in Supp. Fig. 3.

**Table 1:** Method comparison. All methods are compared on synthetic (top) and real WGBS data (bottom). Runtime of alignment and methylation calling (32 threads) and accuracy given as Root Mean Squared Error (RMSE, lower is better) and Spearman correlation (higher is better) are reported. The number of CpGs for which no reads were aligned (#missed CpGs) is given in a separate row.

|  | Metrics |  | Reduced Alphabet Mapper |  |  | Bisulfite Aware Mapper |  |  |
| --- | --- | --- | --- | --- | --- | --- | --- | --- |
|  |  |  | Bismark | BratNova | gemBS | BSmap | FAME | Segemehl |
| Synth. data 25M | Runtime | (h:m:s) | 3:47:53 | 3:24:38 | 0:17:57 | 2:54:22 | 0:11:54 | 2:46:03 |
|  | Performance | RMSE | 0.101 | 0.156 | 0.196 | 0.609 | 0.063 | 0.053 |
|  |  | Spearman | 0.93 | 0.86 | 0.78 | -0.34 | 0.97 | 0.98 |
|  | #missed CpGs |  | 12371 | 44800 | 57319 | 583528 | 5130 | 4444 |
| Real data 437M | Runtime | (h:m:s) | 48:13:18 | 28:46:20 | 10:41:38 | 18:12:25 | 5:09:46 | 42:50:24 |
|  | Performance | RMSE | 0.182 | 0.345 | 0.231 | 0.325 | 0.17 | 0.181 |
|  |  | Spearman | 0.86 | 0.61 | 0.8 | 0.79 | 0.88 | 0.86 |
|  | #missed CpGs |  | 132704 | 747904 | 442318 | 28208 | 88470 | 71571 |

**Figure 4:**
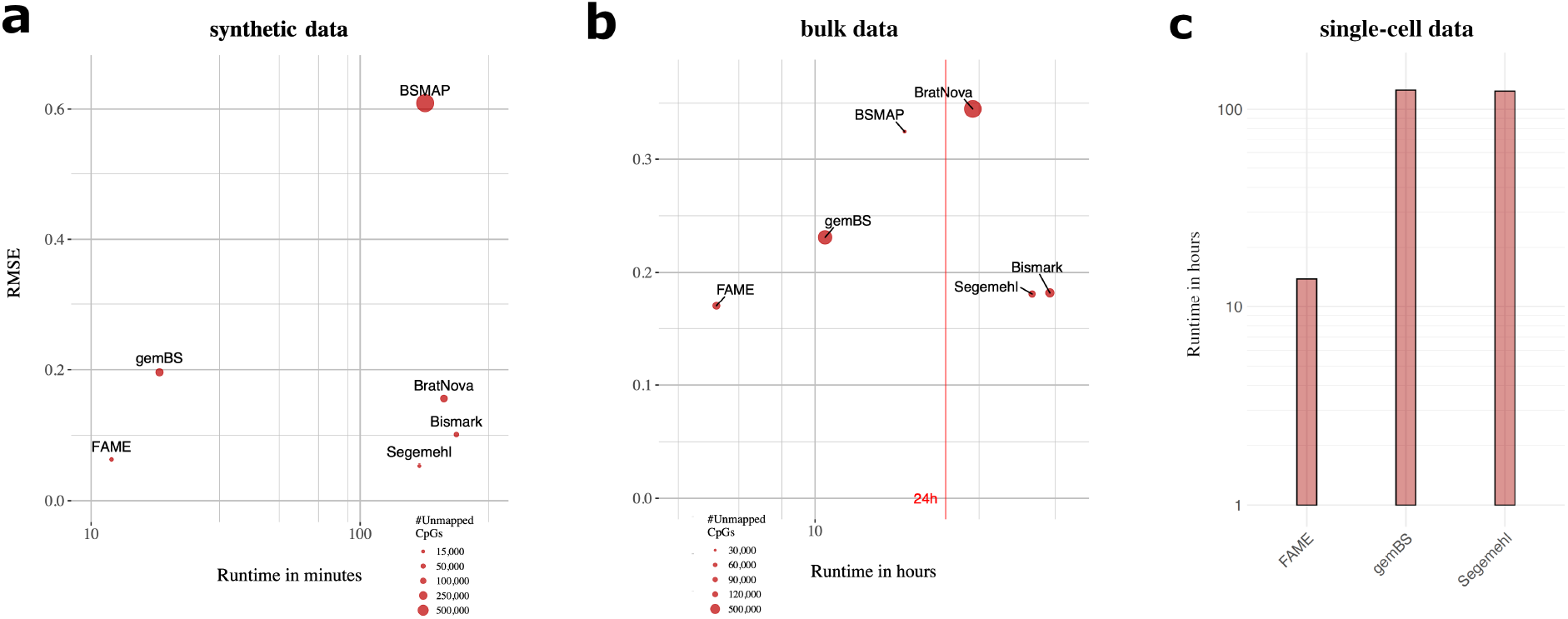
Comparison on bisulfite sequencing data. Visualization of results for simulated bulk sequencing WGBS reads (**a**) and real WGBS data (**b**) of LNCaP cell lines (Pidsley et al. 2016). We compare runtime (x-axis) against error of the predicted methylation values as RMSE (y-axis). The size of each point indicates the number of unmapped CpGs. In case of the real world data, EPIC arrays serve as baseline methylation calls against which we compare the methods. Runtimes in hours on log-scale for scBS-seq (**c**) of 192 cells ((Linker et al. 2019)).

We observe that FAME yields accurate results, improving on all tested RA mappers and yielding as good results as the most accurate mapper Segemehl. More importantly, FAME is running an order of magnitude faster than Segemehl and Bismark, the most accurate RA and BA approaches in our test, bringing down the computation time from several hours to minutes, whereas the second fastest method gemBS showed poor methylation rate calls.

Our results further confirm that BA mappers, such as FAME and Segemehl, yield more accurate methylation rates than RA mappers, which supports the intuition considering the false positive matchings introduced by RA approaches (see Fig. 4c).

### Predicting DNA methylation on real world data against EPIC bead

In the next experiment, we used real WGBS data (437 million paired reads) from the LNCaP cell line, which were accompanied by EPIC array measurements (Pidsley et al. 2016). EPIC arrays provide an alternative large-scale measurement of CpGm values, which is not concerned with the same biases as WGBS.

The results, visualized in Figure 4b and Table 1, show that FAME is 8, 9.5, and 2 times faster than Segemehl, Bismark, and gemBS, respectively. In terms of accuracy, FAME outperforms all RA and BA mappers on this real data set (Figure 4b, Table 1, Supp. Fig. 7, 8). FAME thus provides a unique balance between WGBS processing speed and accuracy. Further analyses showed that Segemehl and FAME are on par with the highest number of covered CpGs, while all other methods show up to an order of magnitude more CpGs with no aligned read. As with the synthetic data experiment, this real data set highlights inferior RA mapper performance compared to BA mappers, which has been independently reported in the literature (Otto, Stadler, and Hoffmann 2012; Frith, Mori, and Asai 2012).

The synthetic data sets used to evaluate the methods and the processed EPIC bead data are deposited online for reproducibility (see Data Availability statement).

### Scaling to single cell data

To this end, the analysis showed that FAME yields accurate results in a fraction of time of the state-of-the-art. The following analysis is concerned with the scalability of methylation calling to the sequencing yields produced from single cell experiments. A unique feature of FAME is to directly support single cell WGBS datasets, where reads of each cell are aligned, counted, and methylation rates are put out per cell, without additional overhead. We thus investigated the performance on single cell data using a dataset of 192 cells from (Linker et al. 2019). We used gemBS and Segemehl to represent the fastest, respectively most accurate, competing method from our previous analyses and applied them on this single cell data set. FAME was able to process all cells within 11 hours, which is an order of magnitude faster than competitors, which both took more than 5 days to finish the analysis (see Figure 4c). All runtimes were achieved while parallelizing on 32 cores on a modern server and demonstrate that single cell data methylation calling is a huge bottleneck, even more with the recent advances of combinatorial indexing and other platforms that yield sequencing of thousands of cells per run (Vitak et al. 2017; Mulqueen et al. 2018).

## Discussion

In the scope of this work, we present a novel method to efficiently quantify DNA methylation from WGBS data and analyze its ability to accurately predict methylation rates on synthetic and real world data in comparison to state-of-the-art methylation callers.

To overcome limitations of evaluations that are solemnly based on mapping efficiency, here we directly evaluate on synthetic data how close the predicted methylation rates are to the ground truth. Thus, any inefficiencies in read mapping, wrongly mapped reads, or mapping biases are captured by this evaluation as they lead to wrong or biased methylation estimates, which otherwise could be missed. While we generated the synthetic data in close resemblance of how reads are actually produced by a sequencing protocol, there might be issues in real world data that are not captured by this data set. We, hence, compared on real-world data with the same underlying idea, to leverage baseline methylation rates to evaluate the estimation capabilities of the methods at hand. The actual methylation rates of a cell are unknown, yet we can do our best to evaluate on real data taking into account orthogonal experiments of quantification of DNA methylation. Here, we leverage estimates from EPIC arrays as comparison to the WGBS calls of each tool. While not being an actual ground truth, it gives a different view on methylation rates that does not suffer from the issues inherent in the process of read alignments with asymmetric mappings (Pidsley et al. 2016). In our analysis we settled for the EPIC bead data, as it is to the best of our knowledge the most reliable and state-of-the-art protocol for genome wide methylation analysis besides WGBS.

Overall, both synthetic and real world evaluation matched our expectations that bisulfite aware mapper yield more accurate methylation estimates than tools that use a reduced alphabet for mapping, which is in line with previous findings (Otto, Stadler, and Hoffmann 2012; Frith, Mori, and Asai 2012). The state-of-the-art bisulfite aware mapper Segemehl yields consistently highly accurate results, but is prohibitively slow in comparison to other tools. The widely used tool Bismark also showed consistently reasonable per-formance despite being a reduced alphabet mapper, yet is less accurate than Segemehl or FAME. To our surprise, the method gemBS showed poor estimation quality in our tests. Similar issues of gemBS have been reported independently in a recent comparison study on plant data (Grehl et al. 2020). Our method FAME shows among the best performance regarding estimation quality, while being an order of magnitude faster than the state-of-the-art. Especially for recent large-scale single cell sequencing data sets, this eliminates the need for high performance computing capabilities, which opens the alignment and methylation calling of WGBS single cell data sets to a broader community.

The focus of our work lies on the estimation of DNA methylation from WGBS data. An interesting avenue for future work would be an adaption of our algorithm to NOME data by building the index for GpCs instead of CpGs. Hence, we can use FAME to get open chromatin calls from NOME-seq experiments (Rhie, Schreiner, and Farnham 2018), leveraging FAME’s accuracy and speed for a different data modality. Similarly, adaptations of our index and algorithm to accommodate SLAM-seq based experiments (Muhar et al. 2018) would make for exciting future work. In SLAM-seq – Thiol(SH)-linked alkylation for the metabolic sequencing of RNA – 4-thiouridine (4sU) labeled mRNAs allow to directly quantify the amount of (labeled) mRNA, as a T¿C conversion is prompted when doing 3’-end mRNA-sequencing on labeled fragments but not on unlabeled fragments. In both application scenarios, NOME-seq and SLAM-seq, correctly resolving the asymmetric mapping is crucial to obtain accurate results and suitable methods are needed for fast processing for these new generation of techniques.

## Conclusion

We considered the problem of efficiently and accurately mapping reads from bulk and single-cell WGBS experiments. Current methods are either slow – imposing bottlenecks on the analysis of the reads – or inaccurate. What all of them have in common is that they are diskspace-intensive with terabytes of data produced as byproducts for a single experiment, posing a challenge for modern single-cell dataset analysis. Apart from the inaccuracies introduced by many alignment tools, this poses a challenge especially in the now common cloud-computing settings, where both runtime as well as diskspace directly translate to costs.

Here, we introduced FAME, a fast and accurate WGBS aligner that is specifically tailored to solve the asymmetric mapping problem introduced by bisulfite treatment. As the approach is fully self-contained and directly models CpGm values in memory, it further avoids unnecessary intermediate alignment files that make the whole process expensive and slow. On both synthetic as well as real data, we show that FAME is as accurate as the best of state-of-the-art methylation caller, yet orders of magnitudes faster and with an order of magnitude less required diskspace, while maintaining a similar memory consumption.

A unique feature of FAME is to directly support single cell WGBS datasets, where reads of each cell are aligned, counted, and methylation rates are output per cell, without additional overhead. We investigated the performance on single cell data (Linker et al. 2019). We used the state-of-the-art gemBS and Segemehl to represent the fastest respectively most accurate competing methods from our previous analysis and applied them on the same single cell data. FAME was able to process the full dataset within 11 hours, whereas both other methods took more than 5 days, all being parallelized on 32 cores.

In conclusion, FAME paves the way for accurate, large-scale CpGm calling and is ideal for cloud computing, because it needs no specialized hardware or access to large file systems. As the number of WGBS experiments increases, e.g., due to advances in single cell measurements, FAME is prepared to address this data deluge. Furthermore, the novel index and corresponding alignment procedure opens up efficient and accurate mapping in different setting of asymmetric mapping in the future, such as required in NOME-seq and SLAM-seq. FAME is open source and licensed under GPL-3.0 under https://github.com/FischerJo/FAME.

## Data Availability

Our generated synthetic data of our performance analysis is publicly available through https://zenodo.org/record/2574694. The bulk WGBS data by (Pidsley et al. 2016) was deposited through accession IDs SRR4238609 up to SRR4238613 in the Sequence Read Archive (SRA) by the authors of that study. (Linker et al. 2019) published their data through project ID PRJEB15062 through the European Nucleotide Archive (https://www.ebi.ac.uk/ena/browser/view/PRJEB15062), the raw reads of the methylation analysis forming the basis for our experiments.

## Footnotes

1 The hash table implementation is a fast open source implementation of the Hopscotch hashmap. https://github.com/Tessil/hopscotch-map

2 http://www.bioinformatics.babraham.ac.uk/projects/trim_galore/

