## Supplementary Material for "Efficiently Quantifying DNA Methylation for Bulk- and Single-cell Bisulfite Data"

January 27, 2023

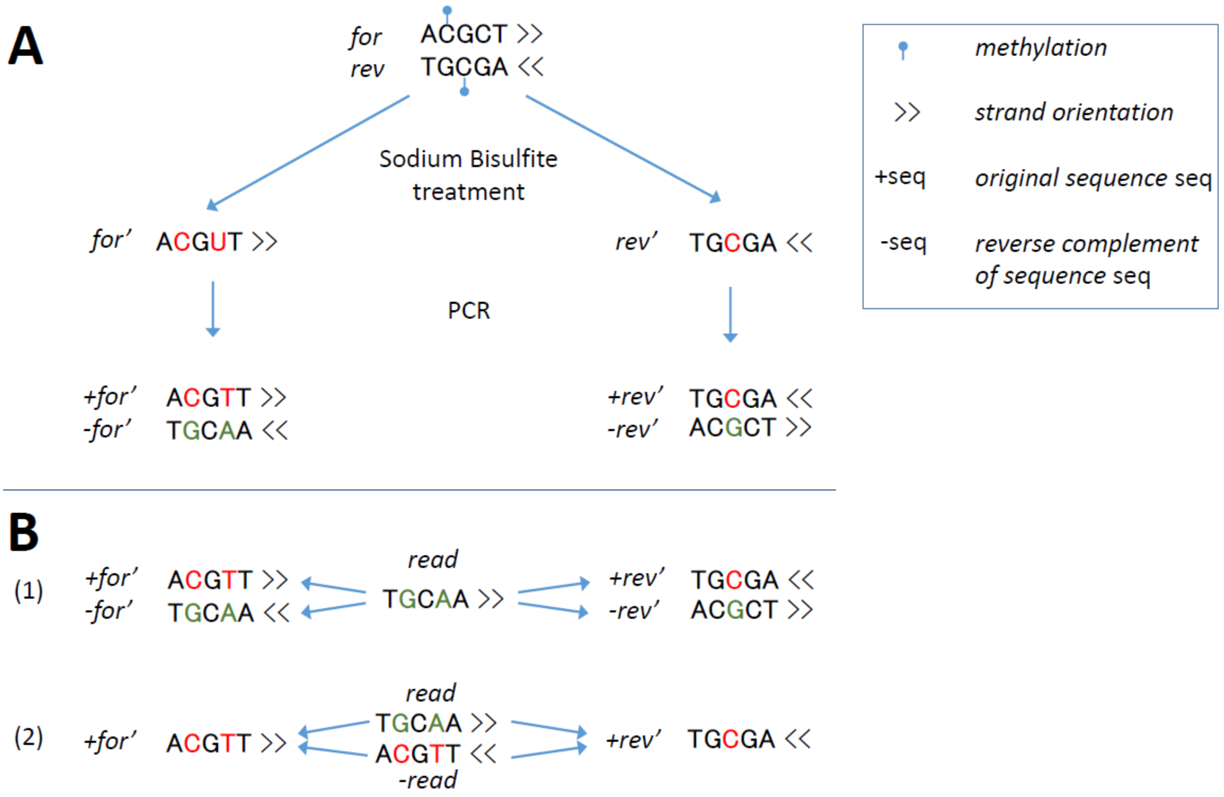


Figure 1: *Reference sequence space blowup in WGBS.* **A**: The two major steps of the WGBS protocol are visualized, leading to a 2-fold blowup of the reference sequence space. First, the reference sequence is denatured and treated with sodium bisulfite, leading to a C to T conversion of unmethylated Cytosines. Positions with Cs in the origin strand (before sodium bisulfite treatment) are consistently colored red. In the PCR step, strand fragments are amplified, leading to sequences representing strand fragments (+) and their reverse complement (-). This results 4 different strands, due to the reverse complementarity of A to T at positions where there was an unmethylated C in the original strand (all positions with former Gs colored in orange). **B**: The read mapping problem in the WGBS sequence space, tackled trivially by mapping to the full reference space (1) and mapped using the idea that either the read itself or its reverse complement must map to the C/T transformed version of the reference (2). The latter circumvents the 2-fold blowup in space requirement for the reference.

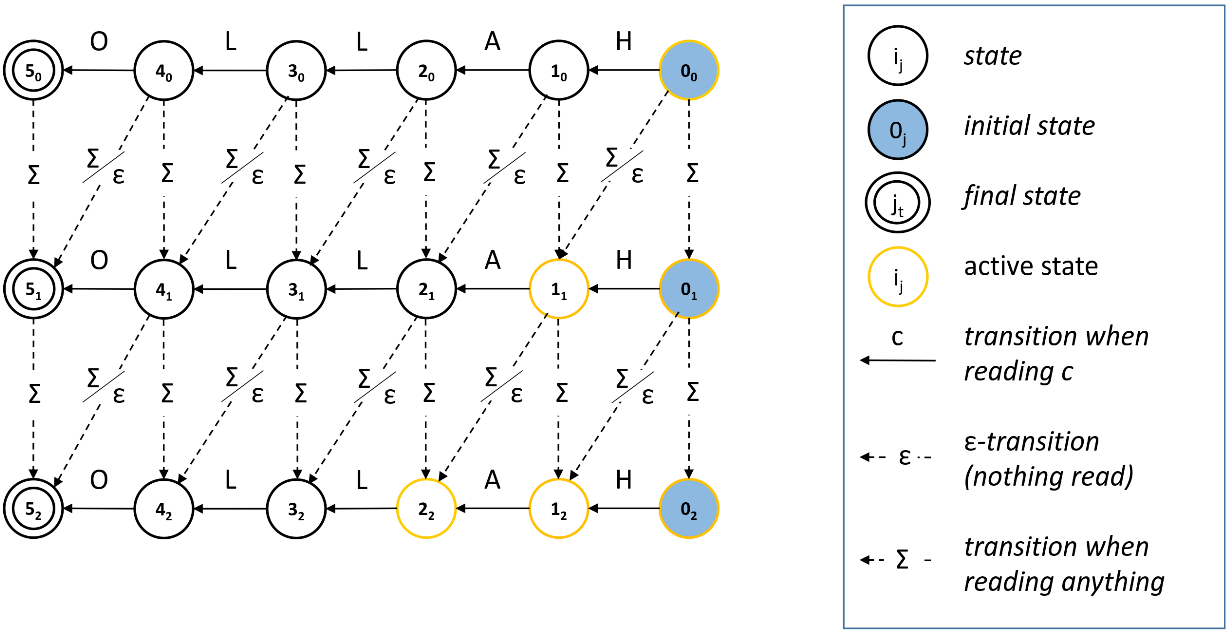


Figure 2: *Approximate shift-and automaton.* An example of an approximate shift-and automaton for the pattern **Hallo** with a maximum of $\tau = 2$ errors. A state $i_j$ appearing in layer j is known to have already produced j errors. The transitions when reading a character c, indicated by full arrows, are just as in the normal shift-and algorithm. Transitions introducing an error in the match will cause a step into the next lower layer. These transitions are indicated by a dashed arrow. $\Sigma$ transitions are triggered when reading any character of the text, corresponding to insertions (vertical transitions) and substitutions (diagonal transitions). The $\epsilon$-transitions can always be performed, without the need of reading a character from the text, corresponding to a deletion.

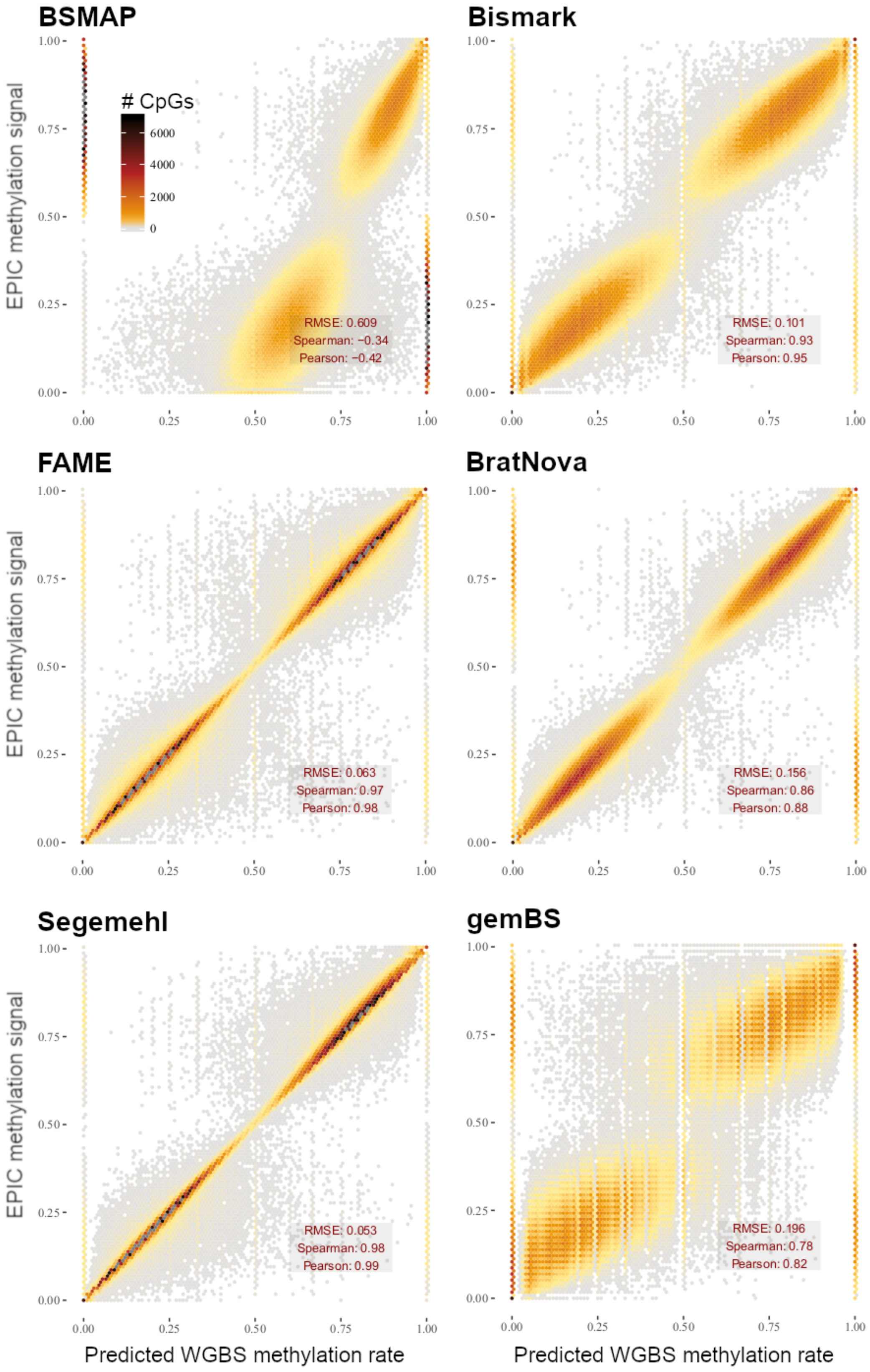


Figure 3: *Synthetic data performance per tool.* The performance of all tools on the synthetic data set is visualized as heated scatterplots with predicted methylation rate on the x-axis and actual methylation rate on the y axis. The color of a point indicates the number of samples (CpGs) falling into this area of the plot. Accuracy metrics are given as RMSE, Spearman, and Pearson correlation coefficient in the bottom right of each plot.

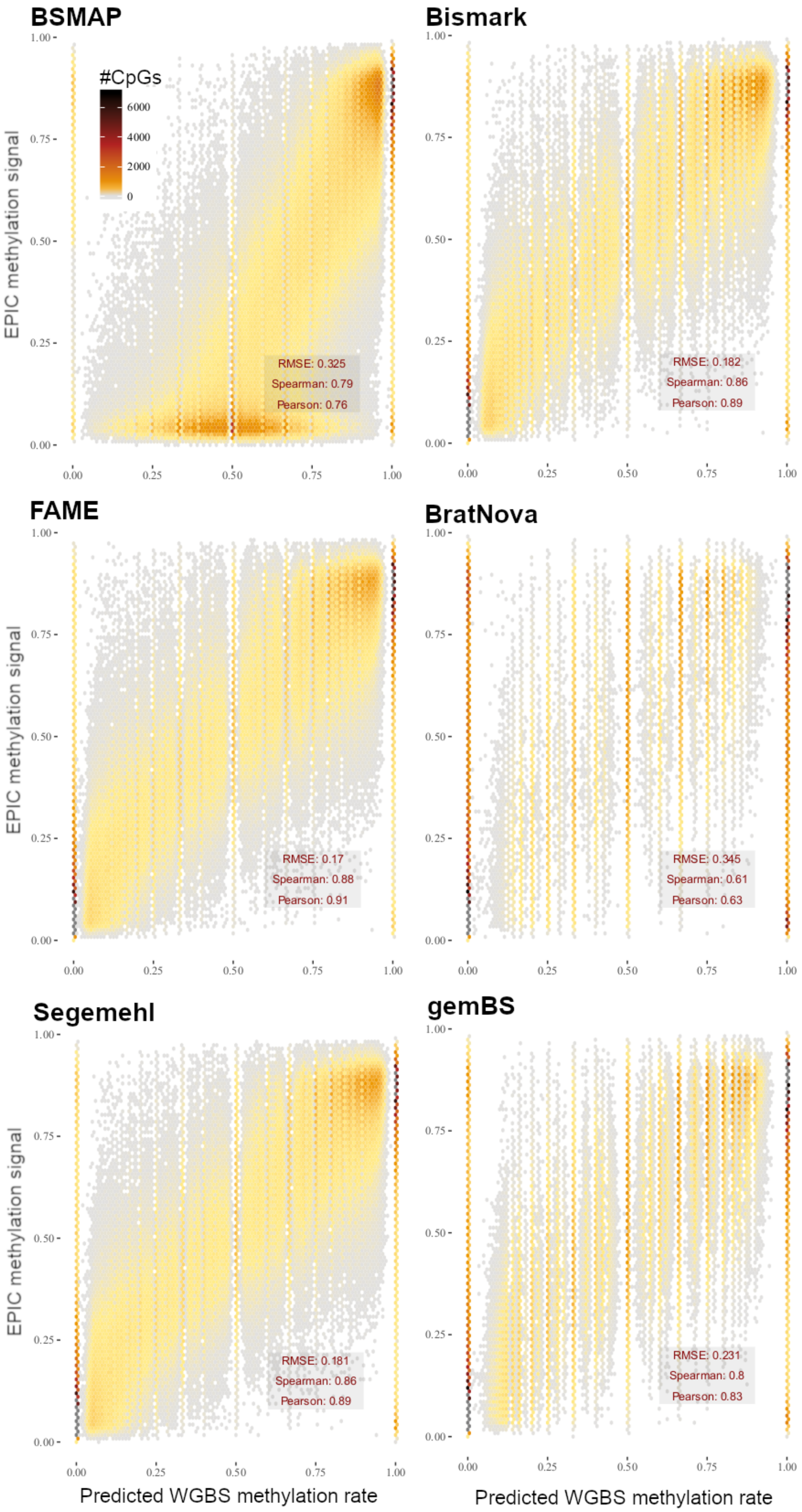


Figure 4: *LNCaP Pidsley data performance per tool.* The performance of all tools on the LNCaP Pidsley data set is visualized as heated scatterplots with predicted methylation rate on the x-axis and actual methylation rate on the y axis. The color of a point indicates the number of samples (CpGs) falling into this area of the plot. Accuracy metrics are given as RMSE, Spearman, and Pearson correlation coefficient in the bottom right of each plot.

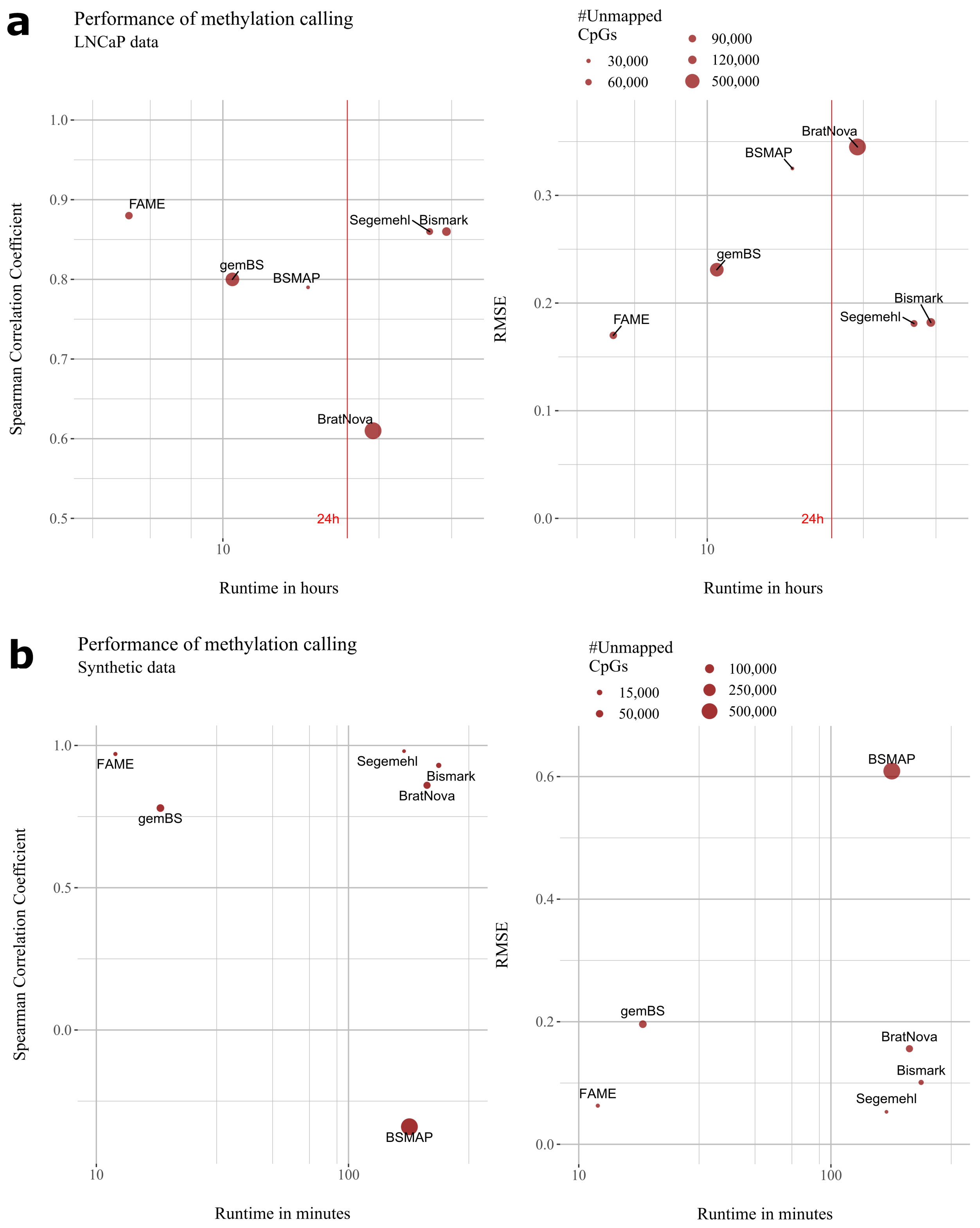


Figure 5: *Performance comparison.* Visualized are the results of all methods measuring runtime (log scale on x-axis) against prediction accuracy (accuracy) in terms of Spearman Correlation Coefficient (left) and RMSE (right). In **a**, results are on the LNCaP data set with correlation and RMSE computed between predicted methylation rate and EPIC bead methylation rate for all CpGs present on the bead. In **b**, results are on synthetic data generated from chromosome 22 of the human genome with known ground truth. Accuracy metrics are computed between predicted and actual methylation rate of all CpGs on chromosome 22.

| Method | Memory in GB |
|---|---|
| BSMAP | 28 |
| BratNova | 8 |
| Bismark | 50 |
| FAME | 35 |
| gemBS | 32 |
| Segemehl | 55 |

Table 1: Rounded numbers for memory consumption in GigaByte (GB) for each tool in the comparison. The RAM consumption was measured on the Pidsley data set.

Table 2: Summary of available tools related to WGBS alignment and methylation calling. We consider different categories: is the software able to call methylation values from alignment result files (Meth. calling)? Does the approach use a BA mapping approach (Bisulfite-aware)? Does the software support non-directional reads , BAM output or input of gzipped reads as well as support of multicore parallelization? The last column indicates whether a tool was included.

| Method | Alignment Method | Meth. calling | Bisulfite-aware | Non-directional | BAM | Gzip | Multi-threading | Included in comparison |
|---|---|---|---|---|---|---|---|---|
| Bat-Meth [1] | FM-index | No | No | Yes | - | - | No | No, too slow in [2] |
| Bat-Meth2 [1] | Batalign | Yes | No | Yes | No | Yes | Yes | No, slower than Batmeth and Bismarck |
| Bismark [3] | bowtie /HISAT | Yes | No | Yes | Yes | Yes | Yes | Yes |
| Bisulfighter [4] | LAST | Yes | Yes | No | - | - | Yes | No, no unstranded PE-read support |
| Brat-BW [5] | FM-index | Yes | No | - | No | - | No | No, predecessor of Brat-NOVA |
| Brat-NOVA [6] | FM-index | Yes | No | Yes | No | No | No | Yes |
| BSMAP [7] | custom | Yes | Yes | Yes | Yes | Yes | Yes | Yes |
| BSSeeker2 [8] | bowtie/SOAP /rmap | Yes | No | No | Yes | No | No | No, too slow in own exp. and [2] |
| BSSeeker3 [9] | SNAP | Yes | No | No | Yes | Yes | - | No. software not working, contacted authors[1] without success |
| BWA-meth [10] | BWA-mem | No | No | Yes | Yes | Yes | Yes | No. Only, aligner. |
| gemBS [11] | GEM | Yes | No | Yes | Yes | Yes | Yes | Yes |
| GSNAP [12] | custom | No | Yes | Yes | No | Yes | Yes | No. Only, aligner. |
| LAST [13] | LAST | No | Yes | No | Yes | Yes | Yes | No. No unstranded PE-read support. |
| WALT [2] | Hash table | No | No | No | No | No | Yes | No. Only, aligner. |
| MAQ [14] | custom | No | Yes | No | No | No | No | No. Outperformed by [3] and [8]. |
| RMAPbs [15] | RMAP | Yes | - | - | No | No | No | No. Outperformed by [3] and [8]. |
| Segemehl [16] | enhanced suffix array | Yes | Yes | Yes | Yes | Yes | Yes | Yes |

[1] The algorithm was not able to align reads and the problem could not be fixed by the authors see github thread: `https://github.com/khuang28jhu/bs3/issues/3`

[2] `https://arxiv.org/abs/1401.1129`

### 1 Computational challenges of WGBS

In Whole Genome Bisulfite Sequencing (WGBS), which is considered the gold standard for methylome analysis of genomes, sodium bisulfite is exploited to induce the conversion of unmethylated Cytosines to Uracil. By subsequent PCR cycles on the genomic fragments, Uracils will be replaced by Thymines, because the Polymerase is insensitive against Uracil and will read it as Thymine [17]. When mapping the reads of the experiment to the reference, a Thymine in the read is allowed to map to a Cytosine in the reference, which means that this Cytosine was unmethylated in the sample.

When carrying out a bisulfite sequencing protocol, the sample DNA is denatured first to allow for the sodium bisulfite treatment, triggering chemical reactions leading to the conversion of all unmethylated Cytosines to Uracil. Then, the two strands are amplified separately using PCR. The subsequent steps are similar to the normal sequencing protocols, including fragmentation of the genome by e.g. sonication or treatment with nucleases, followed by library preparation steps such as adaptor ligation [17]. The two major protocols, considered as standard for WGBS are termed MethylC-seq by Lister et al. and and BS-seq by Crokus et al. An overview of both protocols, which differ in the used adaptor sequences and fragment processing steps, is given in e.g. [18]. Once the experimenter obtains all reads by sequencing the library, the computational post- processing to align the reads to the reference genome has to be performed.

There is one major challenge emerging when it comes to the alignment of WGBS reads in silico. Due to the bisulfite treatment, unmethylated Cytosines (Cs) appear as Thymines (Ts) in the PCR product and Guanines (Gs) appear as Adenines (As) in the reverse complement PCR product. Consequently, we need to allow a match of read Ts to reference Cs, as well as read As to reference Gs (see Fig. 1). However, read Cs mapped to reference Ts and read Gs mapped to reference As are considered as an error in all traditional aligners. We call this the asymmetric mapping problem (depicted in Fig. 1 in main paper), which needs to be accounted for in the alignment phase.

### 2 Related work

There exist a plethora of software packages that deal with WGBS methylation calling that can be classified into two paradigms. Reduced alphabet mapper (RAs) simplify the mapping problem by using two versions of the reference genome, one version with all Cs replaced by Ts and one version with all Gs replaced by As (see Figure 1A). This allows the mapper to leverage the potential of classical alignment tools such as Bowtie [19, 20]. However, this approach comes with two disadvantages, first it introduces false positive matches by ignoring the asymmetric mapping, and second only alignment files are produced, which requires an additional time consuming step to index all reads at a given CpG position and count methylated and unmethylated Cs at this positions. Bisulfite aware (BA) mapper, on the other hand, allow for the asymmetric mapping and hence do not introduce false positive matchings, but can not rely on fast classical alignment software.

The bisulfite aware methods were historically the first developed for WGBS experiments, such as BSMAP [7], RMAP [15], and Segemehl [16]. In our analysis we compare against BSMAP, the fastest BA and more accurate than RMAP, and Segemehl, which shows very accurate methylation calls. We could not include the LAST aligner [13] and its bisulfite enhancement Bisulfighter [4] as they do not support unstranded PE reads (communication with the authors).

In contrast to BA mappers, there exist many RA mappers that are usually wrapper methods for established classical DNA alignment software. Bismark is the most widespread tool for WGBS methylation calling, which is an RA build around alignment software packages such as Bowtie [3, 19, 20]. BSSeeker2 [8] uses an approach for the mapping that is highly similar to Bismark, but also offers processing for Reduced Representation Bisulfite Sequencing experiments. To reflect a real world scenario, we ran all tools on a single server, where in initial experiments BSSeeker2 ran slower than Bismark, which is why we did not incorporate this tool in our comparison. Recently, the new version BSSeeker3 was published that includes a realignment step to account for the asymmetric mapping [9], but we were not able to get the tool running and the authors could not fix the problem (the aligner reported no read alignments[2]. BratNova is a reduced alphabet mapper that uses only one FM index for forward and reverse strand, an extension of the idea of Bismark's usage of several FM indices (one for each strand). This is space efficient and reduces postprocessing time [6]. BratNova comes with no native support for parallelization and we asked the authors to supply a functioning FM construction code (not supplied on the website), which was used then in our benchmark. NovoAlign bisulfite is a software package, in which the bisulfite alignment mode is only commercially available and thus excluded from our experiments as we had no access to it. We chose Bismark as the representative for the accuracy of the RA mappers, as it is a well established and widely used tool.

[2]github issue `https://github.com/khuang28jhu/bs3/issues/3`

#### 2.1 Choice of Methods

Although there is a plethora of methods available, we limited our comparison to a representative set of tools. These tools reflect different indexing and alignment approaches. Although there were other methods available, they were either already proven to be slower than one of the chosen methods, did not run even after contacting the original authors, or were not able to produce an actual call of methylation rates on the given data. We compiled a comprehensive list of available bisulfite tools, excluding methods for color based sequencing. Table 2 includes general information about each tool as well as an explanation in case we did not include a tool in our analysis.

### 3 Results

#### 3.1 Performance on synthetic data

We only considered CpGs that were covered by at least one of the tools with more than 5 reads and penalized any missing methylation call for those considered CPGs by setting the predicted methylation rate to 0 if the true methylation rate >0.5, and to 1 otherwise. An overview of the results comparing runtime with achieved RMSE/Spearman Correlation is visualized in Figure 5a. The performance of each individual tool is visualized in Figure 3.

#### 3.2 Hyperparameter tuning

The index structure that we build introduces several hyperparameter. A hyperparameter of minor importance is the size of a MCpG $m$. A small $m$ means that we hash fewer k-mers for a particular MCpG, it is hence less likely to retrieve a false positive by purely looking at k-mer counts in the candidate retrieval phase. However, we have a larger overall number of MCpGs, increasing the potential number of distinct candidates found for the matching phase. As the candidate counting is a major bottleneck, a too small $m$ hence throttles the performance. For large $m$, the counting phase is cheaper, as we need to count for fewer MCpGs. Since we hash more k-mers per MCpG than for small $m$, it is more likely to get false positive candidates that need to be validated in the matching phase. Thus, it is important to have a balance here. During all tests, we sticked to $m = 2048$, as it generally worked well on our syntethic hyperparameter tuning data (as described in the main paper).

A very important parameter is the strictness $t$ (maximum number of occurrences of a distinct k-mer in the reference) of the index optimization that throws out frequent k-mer sequences, influencing the performance of the algorithm dramatically. Smaller $t$ means a stricter, more lossy filter and thus results in less accurate but much faster versions of the algorithm, because these redundant k-mers often correspond to repetitive regions, which would result in retrieving all repetitive regions as candidates everytime a read contains this k-mer. Smaller $t$ is less sensitive in repetitive regions.

The threshold $q$ corresponds to the minimum number of read k-mers matched in a MCpG such that the read is considered as a candidate. We ran a gridsearch along $t$ and $q$ on synthetic data (see Synthetic data section) to find good default values for these hyperparameter. The results in terms of accuracy and runtime are depicted in Fig. 3 in the main paper. We settled for default values $t = 1500$, $q = 5$, as they show a good tradeoff between runtime and prediction accuracy.

#### 3.3 Performance on real data

For the evaluation we looked at all EPIC bead CpG positions that are mapped by at least one tool with more than 5 reads. We compared the predicted methylation rates of all tools for these positions with the EPIC methylation rate. To penalize uncovered CpGs, we set the predicted methylation rate for each uncovered CpG to 0 if the EPIC methylation rate $> 0.5$, and to 1 otherwise, i.e. the worst prediction that would be possible. The results for each individual tool are visualized in Figure 4, an overall summary comparing runtime and RMSE/Spearman Correlation is depicted in Figure 5. The commands to generate the methylation rates are described in Section 5.

#### 3.4 Memory consumption

We compared the memory consumption in terms of maximum allocated RAM at any time in the steps involved in a pipeline on the Pidsley data set. The index was constructed was constructed for the hg19 reference. In summary, all tools consumed memory within the same order of magnitude, with Segemehl showing the highest amount of allocated RAM. All results are given in Table 1. Note that due to difficulties in measuring the memory for Bismark, we resort to the estimate given by the original authors in the Bismark manual[3].

### 4 FAME manual

FAME is available on GitHub under `https://github.com/FischerJo/FAME`. Through a checkout, the full codebase is downloaded, including a version of gzstream[4], Tessil's hopscotch map[5], Google's sparsehash

[3] Appendix (II) description of the multicore parameter (`https://github.com/FelixKrueger/Bismark/tree/master/Docs`)

[4] `https://www.cs.unc.edu/Research/compgeom/gzstream/`

[5] `https://github.com/Tessil/hopscotch-map`

implementation[6], googletest[7], and a modified version of ntHash [21]. To set general parameters like length of a read, insert size, etc, the user has to modify the file CONST.h as described in the README.md. For all hyperparameters of the algorithm we strongly recommend sticking to the default values.

To run FAME, Google's `sparsehash` library needs to be compiled. To achieve this, run the following command in the top level directory of sparsehash:

```
./configure; make; make DESTDIR=include install
```

To build FAME, type `make` in the top level directory of the FAME repository.

Once the program is compiled, the binary file `FAME` is available. With this binary, the whole algorithm can be used. A helper function is available to guide the user through the parameters, called through `./FAME --help`. The printed manual looks similar to the following:

```


SUMMARY
        This program is designed for the computation of methylation levels
        in (large) mammalian genomes. Please specify the desired values
           for
        the parameters in the file CONST.h, and rebuild the project using
        the provided Makefile (first "make clean" then "make").
        All files are provided as command line arguments by the user.


OPTIONS

Options followed by [.] require an additional argument

        --help
        -h                                  Help

        --genome      [.]                   Specification of a filepath to a
           reference genome
                                            in fasta format.

        -r            [.]                   Specification of a filepath to a
           set of reads in
                                            fastq format. If not specified,
                                               index is built and
                                            saved in file provided via --
                                               store_index.

        -r1, -r2      [.]                   Specification of a filepath to a
           set of reads in
                                            fastq format corresponding to the
                                               first resp. second
                                            read infor paired read set

        --gzip_reads                        Read file specified by -r is
           treated as gzipped
                                            file (.gz file ending).

        --store_index [.]                   Store index in provided file in
           binary format.

        --load_index  [.]                   Load index from provided file.
           Note that all
                                            parameters used to build the index
                                                must be the same
```

[6] https://github.com/sparsehash/sparsehash
[7] https://github.com/google/googletest

```
                                              as used in the current CONST.h.
                                                 This will be checked
                                              while loading.

        --out_basename[.]
        -o            [.]                     Store CpG methylation leves in
           specified filepath,
                                              generating a file with name
                                                 basename_cpg.tsv
                                              Format is specified as header in
                                                 first line of file.

        --no_loss                             Index is constructed losless (
           WARNING: VERY SLOW!)

        --unord_reads                         Disable optimization to find
           stranding of reads.

        --human_opt                           The reference genome is treated as
            GRCH or HG version
                                              of the human genome. Unlocalized
                                                 contigs etc are pruned.


 EXAMPLES

        Setting: Read a reference genome and save index for consecutive
           usages.
         /path/to/FAME --genome /path/to/reference.fasta --store_index
            index.bin

        Setting: Load index from previously stored index, map reads stored
            in .gz format.
         /path/to/FAME --load_index index.bin -r /path/to/reads.fastq.gz
```

A full list of options and hyperparameters can be found in the `README.md` file in the top level directory of FAME and on the github page of FAME.

If a WGBS read set is queried to FAME, the software will produce an output file that contains a 6 column table in tab-separated-value (tsv) format. The column contain for each CpG

```
Chromosome   Position   FwdMeth   FwdUnmeth   RevMeth   RevUnmeth
```

where "Position" is the (0-based) position of the C on the forward strand, and "FwdMeth" and "FwdUnmeth" are the number of methylated Cs and unmethylated Cs mapped to the forward strand CpG, respectively. "RevMeth" and "RevUnmeth" are the corresponding counts for the CpG on the reverse strand.

### 5 Commands

In this section we briefly summarize all program calls we made to generate the results of the individual mapping tools.

#### 5.1 Bismark

Index generation:

```
./bismark_genome_preparation --path_to_bowtie /Path/To/Bowtie/
   /Path/To/hg19_ref/
```

Read mapping:

```
./bismark --multicore 5 -X 1500 --non_directional -o tmp_bis/ --gzip
   --path_to_bowtie /Path/To/Bowtie/ --samtools_path /Path/To/Samtools/
   --genome /Path/To/hg19_ref/ -1 LNCaP_pooled_1_trimmed.fastq.gz -2
   LNCaP_pooled_2_trimmed.fastq.gz
```

Methylation calling:

```
./bismark_methylation_extractor --cytosine_report -p --comprehensive
   --merge_non_CpG --
buffer_size 10G --genome_folder /Path/To/hg19_ref/ --samtools_path
   /Path/To/Samtools/ -o methcall_bis/ --multicore 5
   LNCaP_pooled_1_trimmed_bismark_bt2_pe.bam
```

#### 5.2 BratNova

We used a FM construction code that was provided by the main author after contact by email.
Index generation:

```
./brat_nova/build_bw  -r /Path/To/hg19_ref/referencesHG19.txt  -P
   /Path/To/BratNovaIndex
```

Read mapping on synthetic data:

```
./brat_bw -P /Path/To/BratNovaIndex -1
   paired_hg19_CHR22_poisson0_5_p1.fastq  -2
   paired_hg19_CHR22_poisson0_5_p2.fastq  -a 450 -o
   output_resultsSimData.sam -pe
```

Methylation calling (note that files are changed according to BratNova requirements):

```
./acgt-count -r /Path/To/hg19_ref/referencesHG19.txt -s mappedResults.txt
    -P methylomeLNCAP.txt
```

#### 5.3 BSMAP

Read mapping for LNCaP data:

```
./bsmap -a LNCaP_pooled_1_val_1.fq.gz -b LNCaP_pooled_2_val_2.fq.gz  -d
   /Path/To/hg19_ref/hg19.fa -o LNCAPv6out.sam -p 32 -n 1 -v 6 -x 1800
```

Read mapping for synthetic data:

```
./bsmap -a paired_hg19_CHR22_poisson0_5_p1.fastq  -b
   paired_hg19_CHR22_poisson0_5_p2.fastq -v 10  -d
   /Path/To/hg19_ref/hg19.fa -o SimDataNew.sam -z 64 -p 32 -n 1  -r 0  -x
    650
```

Methylation calling:

```
./methratio.py  -o SimDataNew.bsmap -d /Path/To/hg19_ref/hg19.fa -s
   /Path/To/samtools/ -z -u -p -x CG SimDataNew.sam
```

MethylDackel call for error search:

```
$samto view -bS LNCAPv6out.sam | $samto sort - LNCAPv6sorted.bam
./MethylDackel extract -q 0 -p 1 /Path/To/hg19_ref/hg19.fa
   LNCAPv6sorted.bam
```

#### 5.4 FAME

Index generation:

```
./FAME --genome /Path/To/hg19_ref/hg19.fa --store_index
   index_hg19_spaced_k32_w24_6_m2048_t1500 --human_opt
```

Read mapping and methylation calling for LNCaP (insert size set to 1600, remaining parameters to default):

```
./FAME -r1 LNCaP_pooled_1_trimmed.fastq.gz -r2
   LNCaP_pooled_2_trimmed.fastq.gz --gzip_reads --load_index
   index_hg19_spaced_k32_w24_6_m2048_t1500 -o LNCaP_results
```

Read mapping and methylation calling for synthetic data (all parameters to default):

```
./FAME -r1 paired_hg19_CHR22_poisson0_5_p1_Segemehl.fastq -r2
    paired_hg19_CHR22_poisson0_5_p2_Segemehl.fastq --load_index
    index_hg19_spaced_k32_w24_6_m2048_t1500 -o Synth_results
```

Read mapping and methylation calling for single cell data (read length set to 125, as given by the protocol):

```
./FAME --load_index index_grch38p13_spaced_k32_w28_6_m2048_t1500_mod2
    --gzip_reads --sc_output FAME_meth_per_cell.tsv -o FAME_collapsed
    --sc_input FAME_meta.txt --paired --unord_reads
```

#### 5.5 gemBS

We used gemBS version 3.2.1 in the experiments. gemBS uses two files that are setup before running the method. A conf file and csv file, the first details parameters and location of the reference sequence and fastq files and the index. The second file names the experiments and the read files to be used for it.

Experiment gemBS.conf file:

```
reference = /Path/To/hg19.fa
extra_references = reference/conversion_control.fa.gz
index_dir = indexes

base = .
sequence_dir = .fastq/@SAMPLE
bam_dir = ${base}/mapping/@BARCODE
bcf_dir = ${base}/calls/@BARCODE
extract_dir = ${base}/extract/@BARCODE
report_dir = ${base}/report

project = SimDataChr22
species = Human

threads = 32
jobs = 1
[mapping]

underconversion_sequence = NC_001416.1
overconversion_sequence = NC_001604.1
include IHEC_standard.conf

[calling]
contig_pool_limit = 5000000
```

Experiment gemBS.csv file for simulated data :

```
Barcode,Name,Dataset,File1,File2,Library
barcodeDE,Simdata1,SimDatachr22,paired_hg19_CHR22_poisson0_5_p1.fastq.gz,
paired_hg19_CHR22_poisson0_5_p2.fastq.gz,LBsim
```

Experiment gemBS.csv file for Pidsley data :

```
Barcode,Name,Dataset,File1,File2,Library
lncapBarcode,Reads,LNCAPreads,LNCaP_pooled_1_val_1.fq,
LNCaP_pooled_2_val_2.fq,LBLNCAP
```

Then the same commands for index generation are used:

```
gemBS prepare -c gemBS.conf -t gemBS.csv
gemBS index
```

Read mapping and methylation calling:

```
gemBS map
gemBS call -t 32 -j 8
gemBS extract -B
```

##### 5.6 Segemehl

We used Segemehl version 0.3 for the experiments.

Index generation:

```
./segemehl.x -d /Path/To/hg19_ref/hg19.fa -x hg19.ctidx -y hg19.gaidx -F 2
```

Read mapping for LNCaP data:

```
./segemehl.x -d /Path/To/hg19_ref/hg19.fa -i simSegemehl/hg19.ctidx -j
   simSegemehl/hg19.gaidx -t 32  -q LNCaP_pooled_1_val_1_Segemehl.fq -p
   LNCaP_pooled_2_val_2_Segemehl.fq -D 1  -b -o LNCAPDataD1.bam  -F 2
   --maxpairinsertsize 1800
```

Read mapping for synthetic data:

```
./segemehl.x -d /Path/To/hg19_ref/hg19.fa -i simSegemehl/hg19.ctidx -j
   simSegemehl/hg19.gaidx -t 32  -p
   paired_hg19_CHR22_poisson0_5_p1_Segemehl.fastq  -q
   paired_hg19_CHR22_poisson0_5_p2_Segemehl.fastq -b -D 0  -o
   SimDataD0out.bam  -F 2  --maxpairinsertsize 650
```

Sam related sorting and indexing:

```
sam=/Path/To/samtools
$sam sort -m 10G -@ 32  SimDataD0out.bam SimDataD0sort
$sam index SimDataD0sort.bam
```

Methylation calling:

```
./haarz.x callmethyl -d /Path/To/hg19_ref/hg19.fa -u -b SimDataD0sort.bam
   -t 32 -o CpGCallsSegemehl.vcf
```
